# Biofilm/Persister/Stationary Phase Bacteria Cause More Severe Disease Than Log Phase Bacteria – II Infection with Persister Forms of *Staphylococcus aureus* Causes a Chronic Persistent Skin Infection with More Severe Lesion that Takes Longer to Heal and is not Eradicated by the Current Recommended Treatment in Mice

**DOI:** 10.1101/476465

**Authors:** Rebecca Yee, Yuting Yuan, Cory Brayton, Andreina Tarff Leal, Jie Feng, Wanliang Shi, Ashley Behrens, Ying Zhang

**Affiliations:** Department of Molecular Microbiology and Immunology, Bloomberg School of Public Health, Johns Hopkins University, Baltimore, MD; Department of Comparative Medicine, Johns Hopkins University School of Medicine, Baltimore, MD; The Wilmer Eye Institute, Johns Hopkins University School of Medicine, Baltimore, MD

## Abstract

*Staphylococcus aureus* is an opportunistic pathogen that can cause persistent infections clinically. Treatment for chronic *S*. *aureus* infections ranges from at least one week to several months and such infections are prone to relapse likely due to the presence of persistent forms of bacteria such as persister cells. Persister cells, which are bacterial cells that become dormant under stress conditions, can be isolated *in vitro* but their clinical significance in *in vivo* infections are largely unclear. Here, we evaluated *S*. *aureus* persistent forms using stationary phase cultures and biofilm bacteria (enriched in persisters) in comparison with log phase cultures in terms of their ability to cause disease in a mouse skin infection model. Surprisingly, we found that infection of mice with stationary phase cultures and biofilm bacteria produced a more severe chronic skin infection with more pronounced lesions which took longer to heal than log phase (actively growing) cultures. After two week infection, the bacterial load and skin tissue pathology, as determined by hyperplasia, immune cell infiltration, and crust/lesion formation, of mice infected with the more persistent forms (e.g. stationary phase bacteria and biofilm bacteria) were greater than mice infected with log phase bacteria. Using our persistent infection mouse model, we showed that the clinically recommended treatment for recurrent *S*. *aureus* skin infection, doxycycline + rifampin, was not effective in eradicating the bacteria in the treatment study, despite reducing lesion sizes and pathology in infected mice. Analogous findings were also observed in a *Caenorhabditis elegans* model, where *S*.*aureus* stationary phase cultures caused a greater mortality than log phase culture as early as two days post-infection. Thus, we established a new model for chronic persistent infections using persister bacteria that could serve as a relevant model to evaluate therapeutic options for persistent infections in general. Our findings connect persisters with persistent infections, have implications for understanding disease pathogenesis, and are likely to be broadly valid for other pathogens.

## Introduction

Infections with *Staphylococcus aureus* can lead to chronic and difficult-to-treat infections such as recurrent skin and soft tissue infections, osteomyelitis, endocarditis, prosthetic joint infections and biofilm-related infections found in patients with indwelling devices [1]. Treatments for persistent staphylococcal infections ranges from over one week to over three months and can require a combination of drugs [2]. Patients with osteomyelitis caused by MRSA receiving less than 8 weeks of treatment were 4.8 times more likely to relapse than those receiving longer therapy [3]. In extreme cases for those with prosthetic implants such as in joint implants, debridement procedures are practiced in addition to antibiotic therapy. Yet, even with invasive therapies, the cure rate ranges from 52-80% of the patients [4-6]. Overall, treatment strategies for persistent staphylococcal infections are burdensome for the patient and cannot achieve complete cure in the patient.

First described in 1942, Hobby et al. found that 1% of *S*. *aureus* cells were not killed by penicillin and these were called persister cells [7]. The persister cells were not resistant to penicillin and hence, did not undergo genetic changes; these cells were phenotypic variants that became tolerant to antibiotics [8]. A clinical observation was also made as penicillin failed to clear chronic infections due to the presence of persister cells found in patients [8]. Persister cells can be found to be heavily enriched in stationary phase cultures, drug-stressed cultures, and inside biofilms. Bacteria inside the biofilm grow slowly, are representative of stationary phase bacteria, and can comprise persister cells due to the high cell density, nutrient and oxygen limiting environment inside the biofilm matrix [9].

Many groups have pursued drug discovery research in search for better treatment strategies to kill persisters and biofilms [10]. However, most treatment regimens have not undergone clinical trials for persistent infections. One of the reasons is that there is a lack of chronic infection models of persistent bacterial infections in mice. Recently, it has been shown that more persistent forms of *Borrelia burgdorferi* cause more severe arthritis in a Lyme arthritis model suggesting that severity of disease is determined by the status and quality of inocula bacteria [11]. In this study, we established a chronic skin infection mouse model for *S*. *aureus* using “biofilm seeding” and evaluated drug combinations in clearing the infection in this persistent skin infection model. Here, we show that persistent bacteria found in stationary phase and biofilm bacteria caused a more severe chronic persistent skin infection in mice leading to bigger lesions and increased immunopathology. Using our chronic skin infection model, we show that doxycycline + rifampin, the current clinically recommended drug combination used to treat recurrent skin infections, failed to clear the infection and heal skin lesions in our mouse model. Additionally, we further show that persistent forms of *S*. *aureus* were also more virulent and caused increased mortality in *C*. *elegans*.

## Results

### Establishing a chronic skin infection in mice

To establish a severe, chronic skin infection model in mice, we evaluated the clinical outcomes of mice subcutaneously infected with different forms of *S*. *aureus* (log phase, stationary phase, and biofilm bacteria) (Fig 1). Mice infected with persistent forms (e.g. stationary phase and biofilms) have bigger and more severe lesions than mice infected with actively growing forms (e.g. log phase) (Fig 1A-C). Mice infected with persistent forms also have increased histopathology (Fig 1D-G). Infection with bacteria from biofilms showed crust formation, hyperplasia, immune cell infiltration and focal lesion/abscess formation in tissue histology whereas infection with log phase bacteria only showed low levels of immune cell infiltration. Mice infected with stationary phase bacteria developed skin lesions that required at least one week longer to heal compared to mice infected with log phase bacteria (p<0.001) (Fig 2A). At two weeks post-infection, the average lesion sizes of mice infected with stationary phase bacteria and log phase bacteria was 32 mm^2^ and 11 mm^2^, respectively. Despite inoculation with equivalent amounts of bacteria (10^8^ CFU), mice infected with bacteria from cultures with more persister cells such as stationary phase or biofilms harbored elevated bacterial loads (Fig 2B) recovered from the skin injection sites. Two weeks post-infection, mice infected with biofilm bacteria, the most persistent form tested, harbored at least 10^4^ CFU/gram of tissue more bacteria than mice infected with log phase (p<0.05). Mice infected with more persistent forms also had increased histopathology (Fig 2C) as determined by pathological scores than mice infected with log phase bacteria (p<0.05). Overall, these *in-vivo* findings suggest that infection of mice with persistent forms of bacteria (i.e. stationary phase bacteria and biofilm bacteria) led to a more prolonged persistent infection with more severe lesion formation, than log phase bacteria.

**Figure 1.**
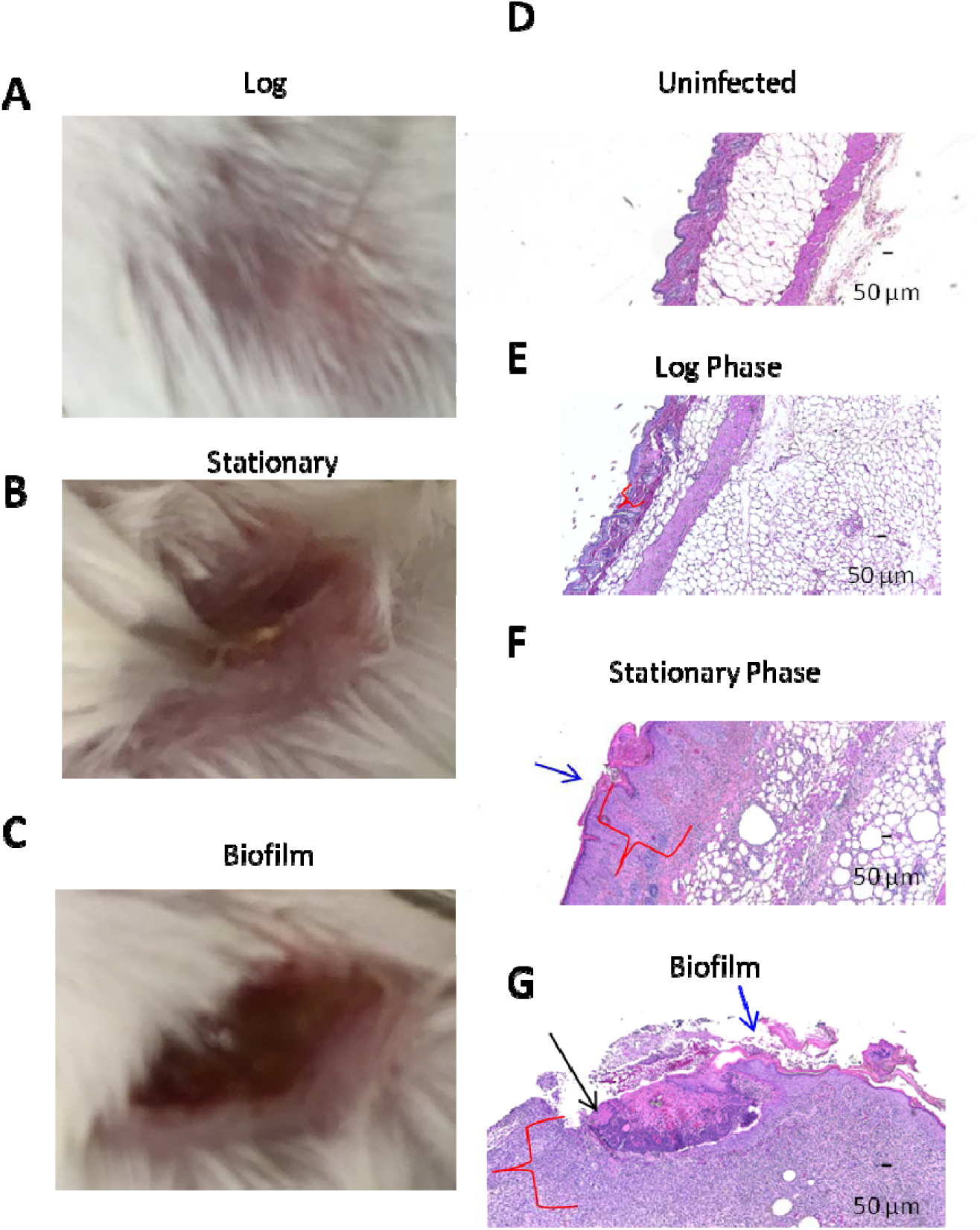
Pathology of mouse skin lesions. Images of gross lesions of infection from log phase bacteria (A), stationary phase bacteria (B), and biofilm bacteria (C). Histopathology of mice with no infection (D), infection with log phase bacteria (E), stationary phase bacteria (F), and biofilm bacteria (G) was performed. Histopathology images were taken at 4X magnification. Blue arrows indicate crust formation, red brackets indicate hyperplasia and cellular infiltration, and black arrow indicate focal lesion development.

**Figure 2.**
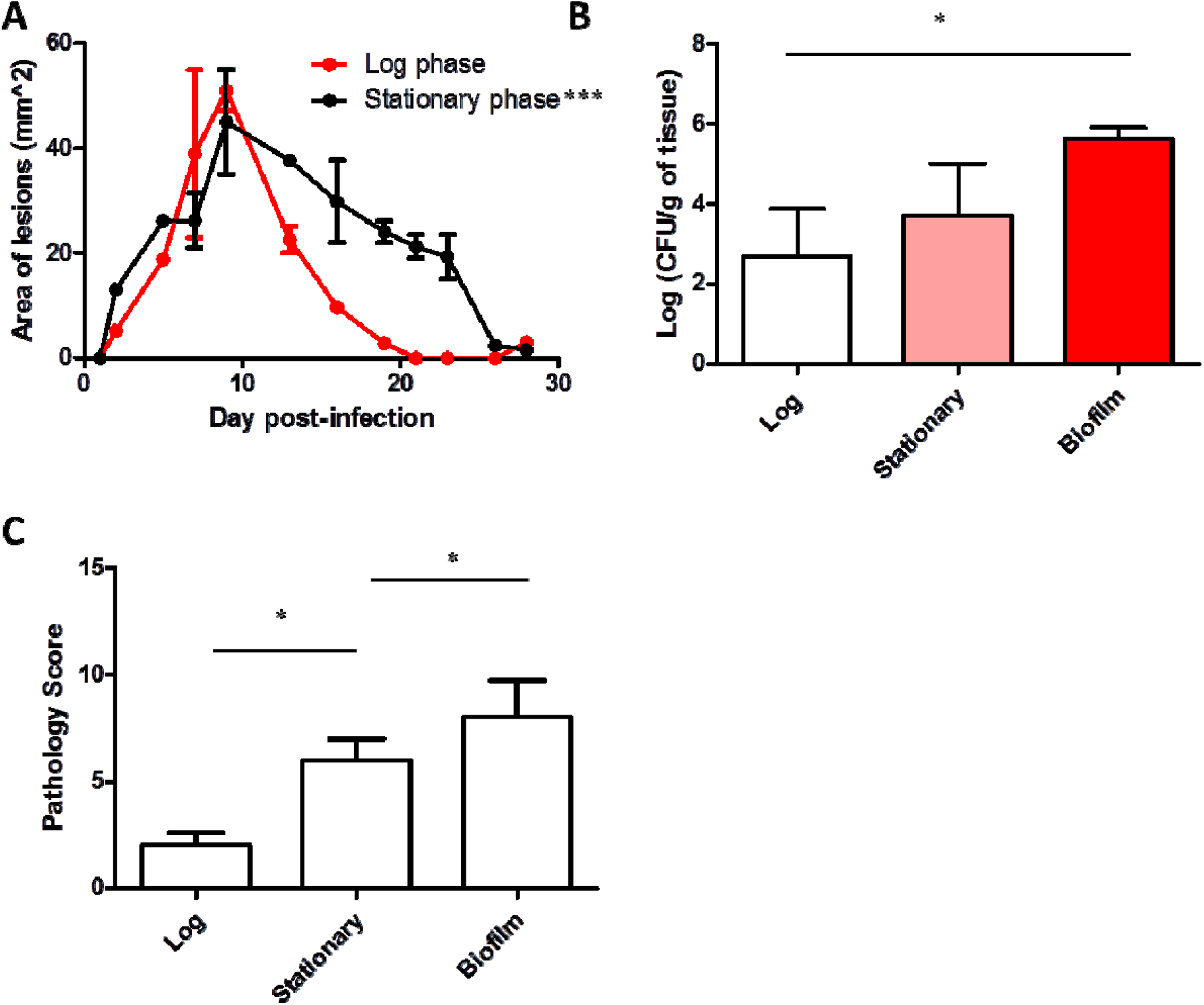
Establishing a chronic *S*. *aureus* persistent skin infection mouse model. The sizes of skin lesions from mice infected with log phase bacteria and stationary phase bacteria were monitored over time (A). After 14 days post-infection, bacterial load (B) was enumerated and pathology scores (C) were evaluated for the skin tissue.

### Clinically used antibiotic treatment failed to clear chronic skin infection in mice

To evaluate the efficacy of treatment strategies in treating the persistent skin infection, we chose to infect mice with bacteria from biofilms of *S*. *aureus* strain USA300 (a MRSA strain) as we observed the most severe clinical presentation (Fig 1 & 2). We allowed the infection to develop for 7 days, followed by treatment of 7 days (Fig 3A). Administration of the combination of doxycycline (50 mg/kg) + rifampin (10 mg/kg), a drug combination used to treat recurrent skin infections[2], did not clear the infection (Fig. 3B). There were no changes in the bacterial load upon treatment. All mice harbored 10^4^ to 10^5^ CFU/gram of tissue. The average change of lesion sizes after treatment for doxycycline + rifampin and no treatment was an average increase of 15% and 200%, respectively, suggesting that while doxycycline + rifampin treatment did not heal the lesions, the treatment, however, did slow the progression of lesion development (Fig 3C). Similar results were seen when mice were infected with *S*. *aureus* Newman strain (a MSSA strain) suggesting that the antibiotic susceptibility profile of the particular strain is not a confounding factor in our results.

**Figure 3.**
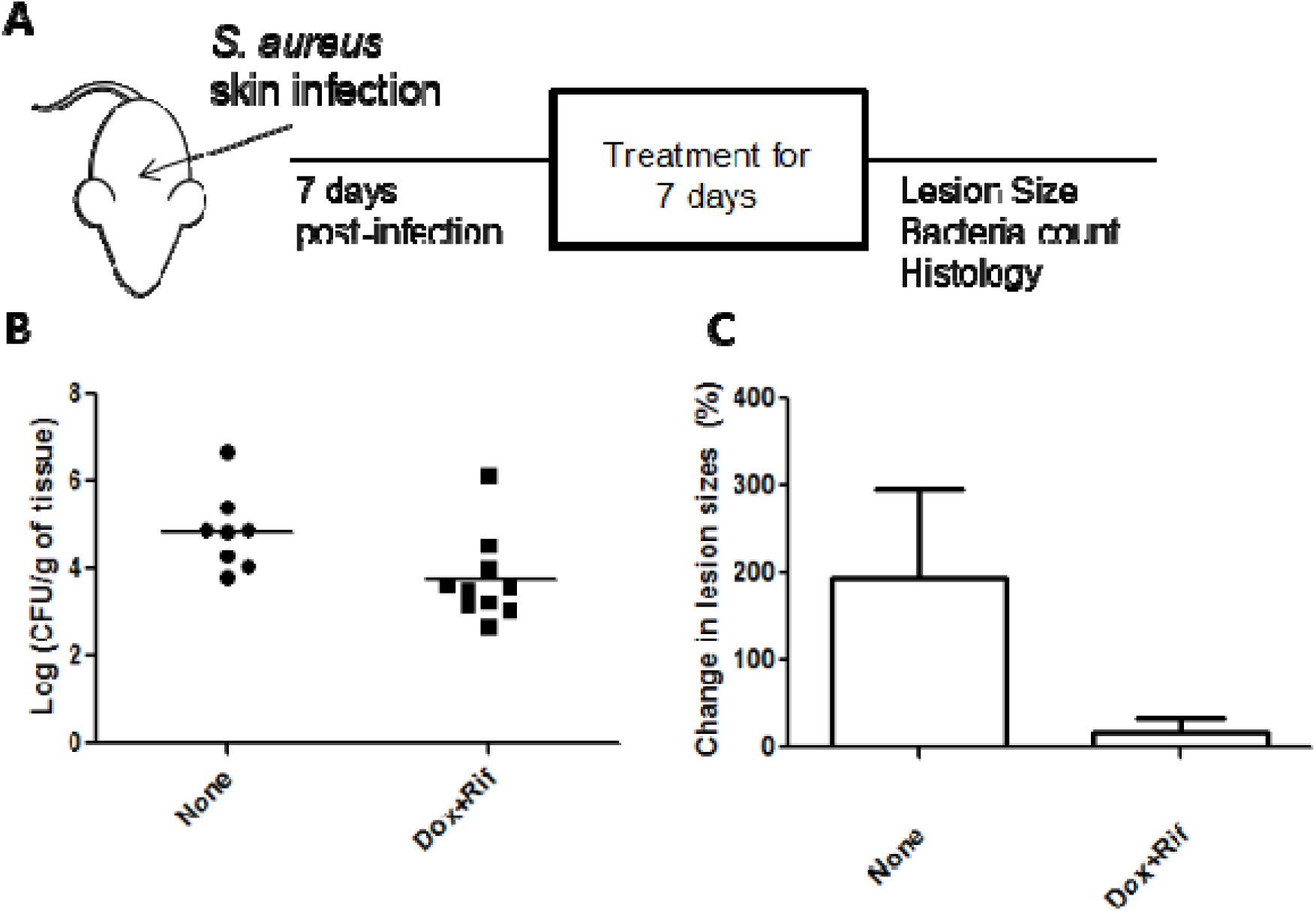
Evaluation of doxycycline + rifampin in chronic skin infection model. Study design of *in vivo* treatment study (A) as described in Methods. Comparison of bacterial load (B) and changes in lesion sizes (C) of mice receiving different treatments.

### Persistent forms of S. aureus caused greater mortality in C. elegans

Besides the clinical outcomes of skin infections, *S*. *aureus* can also cause systemic infection and death in animals such as *C*. *elegans*. We then used a nematode-killing assay to evaluate if persistent forms of bacteria are more virulent and cause more severe disease in *C*. *elegans*. Upon infecting the *C*. *elegans* with different inocula sizes (10^6^ and 10^4^ CFU), the mortality of *C*. *elegans* was the highest in worms that were infected with stationary phase bacteria (p<0.001) (Fig 4). As early as two days post-infection, *C*. *elegans* infected with higher inoculum (10^6^ CFU) of stationary phase bacteria had a survival rate of 60% whereas infection with log phase bacteria resulted in a survival rate of 82% (Fig. 4A). Half of the worms were killed by three days post-infection with stationary phase bacteria but worms infected with log phase bacteria had a survival rate of 78%. Despite the lower inoculum size (i.e. 10^4^ CFU, two log-fold less), the difference in the survival rate of the worms due to inocula status was still observed (Fig 4B). After two days post-infection, the survival rates of worms infected with stationary and log phase bacteria were 64% and 84%, respectively; by day 4 post-infection, the survival rates were 60% and 75%, respectively.

**Figure 4.**
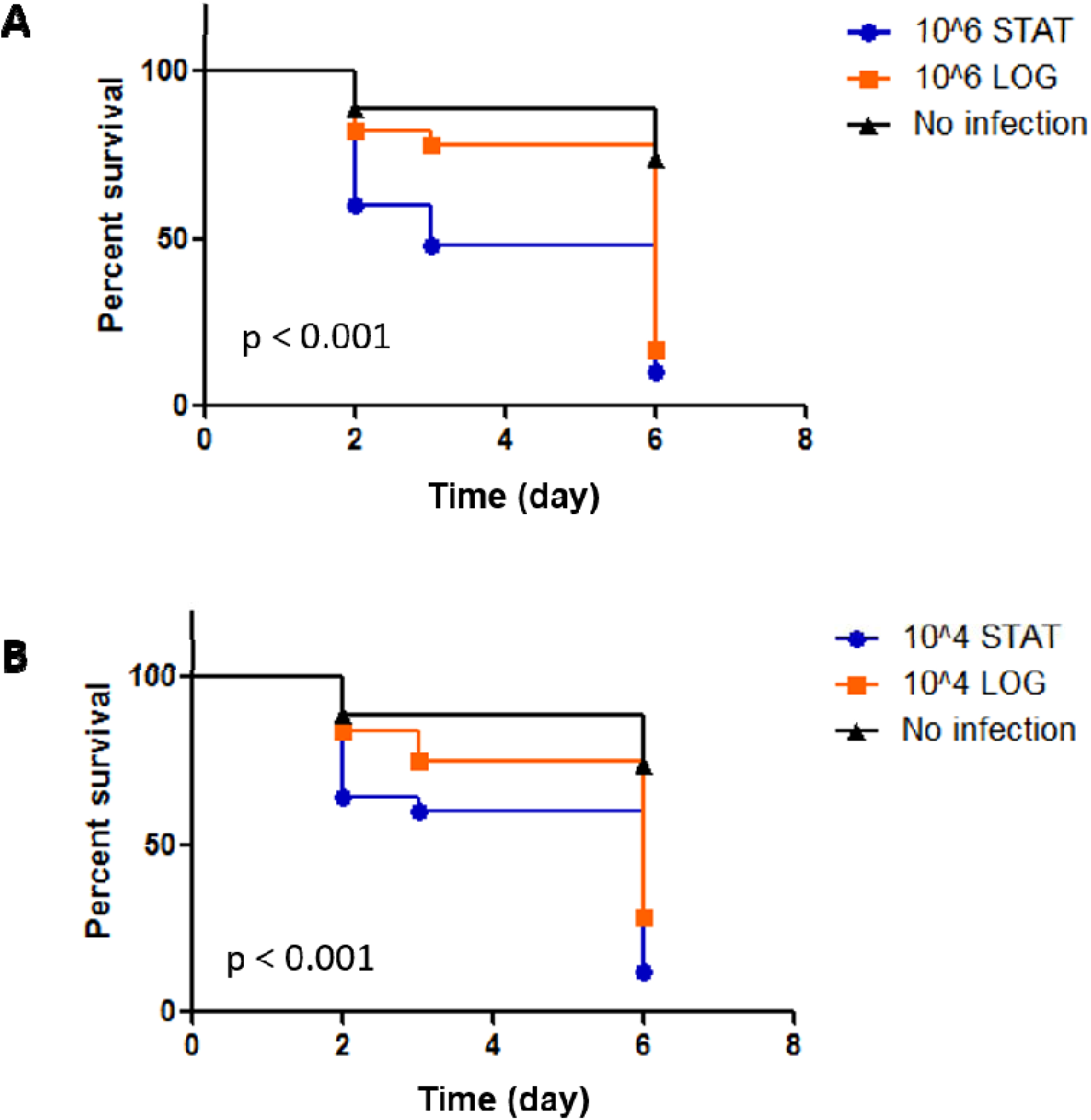
Mortality of *C*. *elegans* infected with different forms of *S*. *aureus*. Infection of *C*. *elegans* with 10^6^ (A) and 10^4^ CFU (B) of stationary phase *S*. *aureus* (STAT) killed *C*. *elegans* faster than log phase bacteria (OG). (Log-rank test)

## Discussion

The discovery of persister cells from Staphylococcal cultures was from many decades ago and it is universally agreed that persister cells and biofilms are hard to kill. The clinical significance of persister cells, which is usually observed in the context of relapsing infections but overall, is generally not well-recognized because the significance of *in vitro* persisters to an *in vivo* clinical phenomenon is unclear. Here, we show that *S. aureus* biofilm bacteria and stationary phase bacteria directly caused a more severe chronic persistent skin infection in mice and formed lesions that were bigger and took longer to heal than log phase bacteria, suggesting a direct clinical consequence of persisters in a host. Treatment of the chronically infected mice with clinically recommended treatment for recurrent skin infection, doxycycline + rifampin, failed to clear the infection and heal the lesions, further showing that current therapies are ineffective against persister bacteria.

It is interesting to note that the bacterial inocula used for animal infections are usually derived from log phase growing bacteria rather than stationary phase or biofilm bacteria [12-15]. The direct association of persistent forms of bacteria causing more severe disease in a host model is an interesting observation that has implications for understanding persistent infection and developing relevant persistent infection models for identifying more effective treatments. Biofilms can be readily found on indwelling catheters retrieved from patients and antibodies against specific biofilm antigens can be isolated from osteoarticular specimens from patients with diagnosed *S*. *aureus* infection in the bone or prosthetic joints [1]. Additionally, recently, using a mouse arthritis model of Lyme Disease, Feng et al. show that infection of mice with biofilm-like microcolony form, a persistent form, of *B*. *burgdorferi* caused the mice to develop more severe swollen joints compared to mice infected with log phase cells [11]. Their findings suggest that “persister-seeding” or “biofilm seeding” can be used to explain why some bacterial infections can become persistent [11]. Our findings here further elaborate that severity of disease is also determined by the status of the inocula bacteria and not completely the bacterial inocula size because our mice developed different disease severity despite the actual bacterial load being comparable. The theory called “Inocula-Dependent Severity of Disease” is not only applicable to *B*. *burgdorferi* but also to common bacterial pathogens such as *S*. *aureus* as shown in this study.

In typical mouse models for *S*. *aureus*, a mouse is infected with a low dose (10^4-6^ CFU) of log phase *S*. *aureus* [12]. In our mouse model, we used a rather high inoculum (10^8^ CFU) and we injected bacteria derived from biofilms. While such conditions can be deemed as artificial, previous animal studies were able to recover a bacterial load of 10^8^ CFU from biofilms formed on heart valves in rabbits with endocarditis, another chronic infection caused by *S*. *aureus* [16]. It is important to note that the chronic infection status of our mice is a key component to our disease model. The phenotype observed in the severity of infection in the host caused by persistent forms cannot be ignored and our model could potentially better mimic chronic infections in humans. While Conlon et al also used a high dose to infect their mice and caused a deep-seated infection, they only allowed the infection to develop for 24 hours before therapy and the mice were made neutropenic [17], a condition that may not apply to a majority of patients suffering from chronic *S*. *aureus* infections as a competent immune system may also induce for persistence.

In conclusion, we show that persistent forms of *S*. *aureus*, such as stationary phase bacteria and biofilm bacteria, can cause chronic persistent skin infections in mice by producing more severe lesions that take longer to heal and have increased pathology than log phase bacteria. Additionally, currently approved regimens are ineffective in clearing these persistent *S*. *aureus* infections caused by these persister forms. Our findings provide a good model for evaluating more effective therapies that eradicate persistent infections in the host. Thus, the power of using a “persister seeding” model [18] to mimic and initiate a chronic persistent infection, has profound implications for understanding disease pathogenesis of many chronic and biofilm infections caused by bacterial, fungal or parasitic pathogens.

## Materials and Methods

### Culture Media, Antibiotics, and Chemicals

*Staphylococcus aureus* strain Newman and USA300 were obtained from American Type Tissue Collections (Manassas, VA, USA). *S*. *aureus* strains were cultivated in tryptic soy broth (TSB) and tryptic soy agar (TSA) from Becton Dickinson (Franklin Lakes, NJ, USA) at 37°C. Doxycycline and rifampin were obtained from Sigma-Aldrich Co. (St. Louis, MO, USA). Stock solutions were prepared in the laboratory,filter-sterilized and used at indicated concentrations.

### Mouse Skin Infection Model and Treatment

Female Swiss-Webster mice of 6 weeks of age were obtained from Charles River. They were housed 3 to 5 per cage under BSL-2 housing conditions. All animal procedures were approved by the Johns Hopkins University Animal Care and Use Committee. *S*. *aureus* strain USA300 and strain Newman were used in the mouse experiments. Mice were anesthetized and then shaved to remove a patch of skin of approximately 3 cm by 2 cm. Bacteria of indicated inoculum size and age were subcutaneously injected into the mice. For log phase inoculum, bacteria grown overnight were diluted 1:100 in TSB and grown for 2 hrs in 37°C at 220 RPM. For stationary phase inoculum, overnight cultures of bacteria grown at 37°C were used. For biofilm inoculum, biofilms were first grown in microtiter plates as described previously [19], and then resuspended and scraped up with a pipette tip. Quantification of all inoculum was performed by serial dilution and plating. Skin lesion sizes were measured at indicated time points up to 4 weeks using a caliper. For the treatment study, treatment was started 1 week post-infection with doxycycline at 100 mg/kg and rifampin at 10 mg/kg twice daily for 7 days by oral gavage as described previously [20]. Mice were euthanized 1 week post-treatment and skin tissues were removed, homogenized, and serial diluted for bacterial plating and counting.

### Histology

Skin tissues were dissected, laid flat, and fixed for 24 hrs with neutral buffered formalin. Tissues were embedded in paraffin, cut into 5-um sections, and mounted on glass slides. Tissue sections were stained with hematoxylin and eosin for histopathological scoring. Tissue sections were evaluated for crust formation, ulcer formation, hyperplasia, inflammation, gross size, and bacterial count and were assigned a score on a 0–3 scale (0 = none, 1 = mild, 2 = moderate, and 3 = severe). The cumulative pathology score represented the sum of each individual pathology parameter. Scoring was performed by an observer in consultation with a boarded veterinary pathologist. Representative images were taken using a Keyence BZ-X710 microscope.

### Nematode-killing Assay

*S*. *aureus* nematode-killing assay was performed as described [21]. *C*. *elegans* N2 Bristol worms (Caenorhabditis Genetics Center) were synchronized to the same growth stage by treatment with alkaline hypochlorite solution as described [22]. Briefly, worms of the adult stage were recovered with M9 buffer and washed twice to remove the residual bacteria in their diet by centrifugation at 1500 rpm for 2 minutes at room temperature. Hypochlorite (5%) was then added and incubated with the worms for 9 minutes to lyse the adult stages and was stopped by addition of M9 buffer. Bleach was removed by centrifugation at 1500 rpm for 1 minute followed by three more washes with M9 buffer. To induce hatching of eggs, M9 buffer was added to the pellet and incubated at 20 °C with gentle agitation and proper aeration. After 24 hours, worms were pelleted with a 2-minute spin at 1500 rpm at room temperature and seeded onto OP50 seeded plates. L4 stage worms were obtained after 48 hours at 20 °C. In each assay, approximately 100 L4-stage nematodes were added to each well containing 1X PBS, 5-Fluoro-2′-deoxyuridine (100 µM) to inhibit progeny development and the indicated inocula of *S*. *aureus*. The plates were incubated at 20 °C and scored for live and dead. A worm was considered dead when it failed to respond to touch and move as observed under a microscope.

### Statistical Analysis

Statistical analyses were performed using two-tailed Student’s *t*-test, two-way ANOVAs, and log-rank comparisons, where appropriate. Mean differences were considered statistically significant if *p* was <0.05. All experiments were performed in triplicates. Analyses were performed using GraphPad Prism and Microsoft Office Excel.

## Acknowledgments

We thank Dr. Jiou Wang for help with the *C*. *elegans* model.

